# Metaplasia Orchestrates Tissue Recovery and Immune Remodeling Through CD73 in Chronic Pancreatitis

**DOI:** 10.64898/2026.11.11.765912

**Authors:** Urvinder Kaur Sardarni, Nicolette Mardik, Victoria A. Gaeth, Brenda Jarvis, Khadija Turabi, Junjun Zheng, Yitian Xu, Xuebo Chen, Jurgen Schrader, Shu-hsai Chen, Benjamin J. Swanson, Jesse L. Cox, Kurt W. Fisher, Holger K. Eltzschig, Richard DiPaolo, Kathleen E. DelGiorno, Jennifer M. Bailey-Lundberg

## Abstract

Chronic inflammatory injury promotes epithelial metaplasia, and progression from metaplasia to dysplasia is a recognized initiating event in carcinogenesis, as metaplastic cells exhibit heightened susceptibility to genetic alterations. In the pancreas, persistent acinar-to-ductal metaplasia (ADM) can progress to pancreatic intraepithelial neoplasia (PanIN), yet the mechanisms that determine whether ADM resolves or persists with fibrosis and sustained inflammation remain poorly defined. Here, we identify CD73 as a central regulator of ADM fate, tissue repair, and immune microenvironment remodeling during chronic pancreatic injury. Analysis of previously published single-cell RNA sequencing dataset from chronic pancreatitis (CP) tissues identified enrichment of Nt5e, which encodes CD73, within Hnf1β⁺ ADM populations. Hnf1bCreERT2 and CD73^fl/fl^;Hnf1bCreERT2 mice were treated with caerulein for 4 weeks to induce pancreatitis and ADM prior to tamoxifen-induced deletion of CD73 in HNF1β-expressing metaplastic epithelial cells. Histologic phenotyping and immune profiling were performed using immunohistochemistry and imaging mass cytometry. Metaplasia-specific CD73 deletion exacerbated pancreatic injury and fibrosis, impaired epithelial proliferation and recovery of the acinar compartment, and altered the abundance and spatial organization of inflammatory cells. Ex vivo ADM assays further demonstrated that CD73 loss increased ADM formation and the production of MCP-1 and TIMP-1. These findings identify CD73 as a context-dependent checkpoint linking epithelial plasticity and immune modulation during chronic pancreatic injury and tissue repair.

## INTRODUCTION

Chronic pancreatitis (CP) is a progressive fibroinflammatory disease characterized by persistent inflammation, progressive fibrosis, exocrine and endocrine dysfunction, and severe pain^1^. In response to chronic injury, pancreatic acinar cells exhibit substantial plasticity and undergo acinar-to-ductal metaplasia (ADM), a protective and initially reversible process that limits tissue damage and necrosis^2^. However, under conditions of sustained inflammation and oncogenic KRAS activation, ADM can persist and progress to pancreatic intraepithelial neoplasia (PanIN), the principal precursor lesion to pancreatic ductal adenocarcinoma (PDAC)^3^. While ADM represents a critical cellular crossroads between regeneration and neoplasia, the molecular mechanisms that govern its resolution versus persistence remain poorly defined.

Inflammatory cells are important regulators of the pancreatic response to injury. Pancreatic injury activates acinar cells and recruits macrophages and neutrophils, leading to the production of pro- inflammatory cytokines and other mediators that shape the local tissue microenvironment. Innate immune cells contribute to the clearance of damaged tissue and support tissue recovery^4^. However, prolonged or dysregulated inflammatory responses can stabilize ADM and promote fibrotic remodeling^5^. Macrophage- derived inflammatory signals have been shown to directly induce ADM, demonstrating that immune cells actively regulate acinar-cell plasticity in response to tissue damage^6,7^. Interactions between myeloid and acinar cells have also been shown to influence acinar-cell survival, redifferentiation and stromal remodeling^8^. Therefore, communication between metaplastic epithelium and immune cells may help determine whether pancreatic injury resolves or progresses toward persistent metaplasia and fibrosis. However, the epithelial factors that regulate this communication and shape the local immune microenvironment during chronic inflammation remain unknown.

Using integrated transcriptomic, histologic, and proteomic profiling of murine and human CP, we previously identified robust induction of NT5E, which encodes CD73, in injury- and KRAS-driven ADM. CD73 is a cell-surface ectonucleotidase that catalyzes the final step in extracellular adenosine production^9–12^. During tissue injury, damaged cells release extracellular ATP, which promotes innate and adaptive immune responses through purinergic receptors. Extracellular ATP is subsequently hydrolyzed to AMP by CD39 and then converted to adenosine by CD73. In contrast to ATP, adenosine generally limits immune activation and protects tissues from excessive inflammatory damage^13,14^. Thus, CD73 serves as an important regulator of the transition from ATP-driven inflammation to adenosine-mediated immune suppression and tissue protection. The consequences of CD73 activity, however, depend on the disease context. Our previous work showed that global loss of CD73 during pancreatic injury resulted in sustained inflammation and metaplasia, indicating that CD73-generated adenosine is important for the resolution of pancreatic tissue injury^15^. In pancreatic cancer, however, CD73-mediated adenosine production is commonly associated with immunosuppression^16,17^. CD73 is also expressed by pancreatic stromal cells, particularly stellate cells and has been associated with fibrosis^18^. Since CD73 is expressed by epithelial, immune, endothelial and stromal populations^14,19^, global deletion could not define its epithelial-intrinsic role in chronic pancreatic injury and metaplasia.

Here, we investigated the functional consequences of genetic deletion of CD73 specifically within metaplastic ducts, focusing on its role in regulating CP severity, fibrosis, and immune composition. We first used acinar-lineage tracing and organoid models to establish that CD73 is induced in acinar-derived ADM following pancreatic injury. We then examined the functional role of metaplasia-associated CD73 using Hnf1bCreERT2 and CD73^fl/fl^;Hnf1bCreERT2 mice. Chronic pancreatitis was induced in these mice by caerulein treatment, allowing ADM to develop before tamoxifen administration to delete CD73 in Hnf1b- expressing metaplastic epithelial cells. This temporally controlled approach allowed us to examine the function of metaplasia-associated CD73 after ADM formation. By combining genetically engineered mouse models with histologic analysis, immune profiling, ex vivo acinar-cell culture and exploratory spatial analysis using imaging mass cytometry, we identified a previously unrecognized role for epithelial CD73 in pancreatic tissue recovery. Loss of epithelial CD73 reduced epithelial proliferation and recovery of the acinar compartment, increased persistent ADM and fibrosis, and altered the inflammatory microenvironment, with increased neutrophil and NK-cell populations and reduced macrophage populations. Together, these findings identify epithelial CD73 as a critical checkpoint that governs whether chronic pancreatic injury resolves or progresses toward persistent metaplasia and fibrosis.

## METHODS

### Mouse Models

All animal experiments involving Hnf1b-CreERT2 mice were approved and performed under the guidelines of the Center for Laboratory Animal Medicine and Care (CLAMC) at the University of Texas Health Science Center at Houston (AWC-24-0011). To genetically delete CD73 from metaplastic ducts that express Hnf1b, the inducible Hnf1b-CreERT2 model was crossed with a CD73^fl/fl^ transgenic mouse (a collaborative gift from Dr. Jurgen Schrader and Dr. Holger Eltzschig) (CD73^fl/fl^) to generate CD73^fl/fl^;Hnf1bCreERT2 mice. All mice were genotyped by tail DNA PCR to confirm their genotypes. Control Hnf1bCreERT2 mice (n=7) and CD73^fl/fl^;Hnf1bCreERT2 mice (n=5), aged 10-12 weeks, were injected intraperitoneally with 5.5 µg of caerulein twice a day, every weekday, for four weeks as previously described^20^. On the final day of caerulein treatment, mice were injected intraperitoneally with 5 mg of tamoxifen. One week after tamoxifen injection, mice were euthanized via isoflurane overdose, and the pancreas was harvested. Pathological analyses and immune profiling were conducted using immunohistochemistry (IHC) and imaging mass cytometry (IMC). Both sexes were used for each experiment, and littermates were enrolled into groups after genotyping.

For acinar-cell lineage tracing, Ptf1aCre^ERTM/+^;LSL-Rosa^eYFP/+^ mice were housed in accordance with National Institutes of Health guidelines in American Association for Accreditation of Laboratory Animal Care- accredited facilities at Vanderbilt University. Lineage tracing was performed as previously described^20,21^. In this model, tamoxifen treatment induced Cre recombinase activity specifically in PTF1A+ acinar cells, thereby initiating expression of enhanced yellow fluorescent protein (EYFP) in these cells. Tamoxifen (5 mg/day, 5 days/week for two weeks) was administered in corn oil by oral gavage to label acinar cells as EYFP+. Chronic pancreatitis was induced with caerulein administered intraperitoneally (IP) at 250 μg/kg, twice per day, for five days/week for two weeks; mice were then allowed to recover for two days before tissue collection. Control mice received PBS.

### Immunohistochemistry and ImageJ Quantification

Tissues were fixed in zinc-buffered formalin, processed, and embedded in paraffin. Sections were placed on positively charged slides and baked at 60°C for 30 minutes. Slides were deparaffinized in histoclear and gradually rehydrated to PBS. Heat-mediated antigen retrieval was performed using a pH 6 solution (Vector Laboratories, H-3300) or a pH 9 solution (Abcam 100x Tris-EDTA, ab 93684). Sections were blocked for 1 hour at room temperature (RT) in 10% FBS in PBST. Primary antibodies were diluted in blocking solution and incubated overnight at 4°C. Primary antibodies used, their dilutions, and antigen retrieval pH are as follows: Murine CD73 (Cell Signaling, D7F9A, 1:200 dilution, pH 6), Human CD73 (Abcam, ab133582, 1:200 dilution, pH6), α-SMA (Abcam, ab5694, 1:500, pH 6), PCNA (Abcam, ab29, 1:400, pH6), Ki67 (Abcam, ab15580, 1:200, pH 6), α-amylase (Abcam, ab199132, 1:50, pH 9), Cytokeratin-19 (Abcam, ab52625, 1:150, pH 9), NIMPR14 (Abcam, Ab2557, 1:100, pH 9), MPO (Abcam, Ab208670, 1:300, pH 9), CD68 (Abcam, ab283654, 1:200, pH 9), CD163 (Abcam, ab182422, 1:500, pH9), CCR2 (Abcam, ab273050, 1:1000, pH9), NK1.1 (Cell Signaling, E6Y9G, 1:50, pH9), TIMP-1 (Abcam, ab216432, 1:250, pH6). Tissues were washed three times with PBS for 5 minutes each and incubated with the appropriate secondary antibody (Vector Laboratories, BA-1000, BA-9400) at RT for 30 minutes at a 1:500 dilution in blocking buffer. Tissues were washed three times with PBS for 5 minutes each and incubated with the Vectastain ABC kit Peroxidase Standard (Vector Laboratories, PK-4000). Tissues were washed three times with PBS for 5 minutes each, and staining was developed using the DAB Peroxidase (HRP) Substrate kit (Vector Laboratories, SK-4100). Tissues were rinsed in tap water for 2 changes of 5 minutes, counterstained with hematoxylin and then gradually dehydrated in ethanol. Tissues were cleared in histoclear and mounted. Analysis of immunohistochemistry images was performed using ImageJ software (https://imagej.net/). 5 to 10 representative fields per tissue were quantified blinded to genotype, depending on tissue size. A freehand selection tool was used to determine the pancreatic tissue area, and the color threshold tool was used to determine positive staining in each field.

### Hematoxylin and Eosin Staining and Analysis

Tissues were placed in histoclear for 8 minutes, 100% ethanol for 4 minutes, and 95% ethanol for 2 minutes. Tissues were then incubated in tap water, hematoxylin and tap water for 3 minutes. Next, tissues were incubated in clarifier, tap water, bluing agent, water, 95% ethanol, eosin, and 95% ethanol for 1 minute each. Tissues were placed in 100% ethanol for 4 minutes and histoclear for 4 minutes, then mounted. The method was performed as previously described^22^. Interstitial edema was quantified as the percentage of edema-positive area relative to the total pancreatic tissue area using Image J. For each tissue section, 5–10 representative fields were quantified.

### Trichrome Staining

Tissues were stained according to the recommended protocol of the trichrome staining kit (ab150686). Tissues were deparaffinized for 9 minutes in histoclear and hydrated gradually in ethanol to distilled water. Slides were placed in preheated Bouin’s fluid for 60 minutes and rinsed under running tap water until the sections were clear. Slides were placed in tap water for 4 minutes and stained with working Weigert’s iron hematoxylin for 5 minutes. Slides were rinsed for 2 minutes under running tap water and then stained with biebrich scarlet/acid fuchsin solution for 15 minutes. Slides were rinsed in distilled water and stained with phosphomolybdic/phosphotungstic acid solution for 10 minutes. Without rinsing, slides were stained with aniline blue solution for 10 minutes. Slides were rinsed in distilled water and then placed in acetic acid solution for 5 minutes. The slides were then dehydrated in 95% ethanol, 100% ethanol and mounted. The method was performed as described previously^15,23^. Collagen deposition was quantified using the color deconvolution plugin in ImageJ. Fibrosis was calculated as the percentage of collagen-positive area relative to the total pancreatic area. For each tissue section, 5–10 representative fields were quantified.

### Immunofluorescence Staining and Quantification

Tissues were fixed overnight in 10% neutral-buffered formalin, embedded in paraffin, sectioned (5 μm thickness), mounted, and stained. Sections were deparaffinized in xylenes, rehydrated in a series of ethanols, and then permeabilized with 0.1% Tween-20 in 1x PBS. Endogenous peroxidase activity was blocked with a 1:50 solution of 30% H2O2:PBS, followed by microwave antigen retrieval in 100 mM sodium citrate, pH 6.0. Tissues were blocked with 5% normal donkey serum and 1% BSA in 1x PBS for 1 hour at room temperature in a humidity chamber. Primary antibodies were diluted in 1% BSA and 0.1% Triton X-100 in 1x PBS and incubated overnight at room temperature in a humidity chamber (Anti-GFP, Abcam, Ab6673, 1:500; TROMA-I, DSHB, supernatant 1.0 mL, 1:200; NT5E/CD73, CST, 13160S, 1:100). Sections were then washed 3 times for 15 min each in 0.1% Triton X-100 in 1x PBS, followed by incubation with Alexa Fluor secondary antibodies (Invitrogen), washed again 3 times for 5 min each in 0.1% Triton X-100 in 1x PBS, then rinsed with distilled water. Samples were mounted with Prolong Gold antifade with DAPI (Invitrogen). Slides were scanned using an Olympus Slideview VS200. Generated .vsi images were imported into QuPath v0.7.0^24^ for analysis. Pancreas tissue was manually annotated, removing islets, lymph nodes, and folded tissue. Pixel classifier was used to classify acinar-lineage (YFP+) region on a per-section basis, generating an acinar annotation for each section. InstanSeg was used to segment cells in the acinar annotation^25^. Then Qupath’s object classifier was used to define CD73 positivity for each cell. Data was exported to .csv for statistical analysis in GraphPad Prism v11.1.0.

### Organoid Generation

Organoids were established from the pancreata of Ptf1aCre^ERTM/+^;LSL-ROSA^eYFP/+^ mice following induction of chronic pancreatitis with caerulein or vehicle control (PBS). The organoid generation protocol was adapted from previously described methods^26,27^. Briefly, pancreata were dissected, minced in Hank’s Balanced Salt Solution (without calcium, magnesium or phenol red) (HBSS) and incubated in digestion media containing 1 mg/mL collagenase I, 10 μg/mL of DNAseI, 1 mg/mL of soybean trypsin inhibitor, and 5ml of dispase diluted in Advanced DMEM/F12 at 37°C for 25-40 minutes, with manual pipetting of pancreata every 5 minutes, until tissue was digested. Then, digestion was quenched with organoid wash buffer containing HBSS (without calcium, magnesium or phenol red), supplemented with 10% FBS, 1% antibiotic-antimycotic (A/A), and 1% L-glutamine. Cells were placed on ice. GFP+ cells were manually picked from the solution using a Leica M205 FA Fluorescence stereo microscope. After picking, cells were filtered through a 70 μm strainer and pelleted at 500 x g for 5 min at 4°C. The pellet was washed by resuspending in organoid wash buffer, and then re-pelleted at 500 x g for 5 min at 4°C. The pellet was embedded in Matrigel and organoid media was added. Pancreatic organoid media consisted of Advanced DMEM/F12 supplemented with 10% Noggin conditioned media, 10% R-spondin conditioned media, 1 x B27, 10 mM nicotinamide, 1.25 mM N-acetylcysteine, 50 ng/mL of mEGF, 100 ng/mL of mFGF-10, 10 nM hGastrin I, and 500 nM of A83-01. Y-27632 was added to media directly after isolation but removed in following passages. Organoids were maintained in 5% CO_2_ at 37°C and passaged at confluency. Organoids were imaged using an ECHO REVOLVE microscope with the Trans, FITC, and Cy5 channels.

### RNA extraction and qRT-PCR

Total RNA was isolated using TRIzol™ Reagent (Thermo Fisher Scientific) according to the manufacturer’s instructions, with few modifications. Briefly, culture media was removed, and samples were washed with ice-cold 1 x PBS. Samples were lysed in TRIzol™ Reagent (300µL/Matrigel dome), vigorously shaken for 15 sec, and incubated at room temperature for 5 min. Chloroform (200µL) was added to each sample, followed by vigorous shaking and incubation at room temperature for 5 min. Samples were centrifuged at 12,000 × g for 15 min at 4°C. The aqueous phase was transferred to a fresh tube and mixed with an equal volume of isopropanol and incubated at room temperature for 10 min before centrifugation at 12,000 × g for 10 min at 4°C. The supernatant was discarded, and the RNA pellet was washed with 75% ethanol (molecular biology grade) and centrifuged at 7,500 × g for 5 minutes at room temperature. The ethanol wash was repeated once. RNA pellets were air-dried at room temperature for 10 minutes with the tube lids open and resuspended in 20 µL of diethyl pyrocarbonate (DEPC)-treated water. Samples were incubated at 60°C for 10 minutes to facilitate RNA dissolution. RNA concentration was measured using NanoDrop 2000.

Reverse transcription quantitative PCR (RT-qPCR) was performed using the Luna Universal One-Step RT- qPCR Kit (E3005, New England Biolabs) on a Bio-Rad CFX96 Real-Time PCR Detection System. Data were analyzed using CFX Manager 3.1, using Rplp0 as a reference gene. The primer sequences (5′–3′) were as follows: Nt5e, forward: GCAAACATTAAGGCACGGGG and reverse: TCACCGCCAACAGAGAGAAC; *Rplp0,* forward: GCCAATAAGGTGCCAGCTG and reverse: CTCCCACCTTGTCTCCAGTC.

### Analysis of Published RNA Sequencing Datasets

Published single cell RNA sequencing (scRNA-seq) data were analyzed as previously described^15^. Briefly, processed count matrices for the scRNA-seq datasets from Ma et al.^20^ were downloaded from the Gene Expression Omnibus (GEO) database (accession number GSE172380). Low-quality cells were filtered based on read counts, the number of genes expressed, and the proportion of mitochondrial reads using the thresholds described in the original publication. Filtered gene count matrices were log-normalized, and the top 2,000 variable features were scaled prior to principal component analysis (PCA) and uniform manifold approximation and projection (UMAP) using the R package Seurat^28^. Seurat cell clusters were annotated using marker genes reported by Ma et al^20^.

### Imaging Mass Cytometry and Data Analysis

Imaging Mass Cytometry (IMC) enables the simultaneous detection of 22 protein markers for in situ single-cell categorization. One mouse per genotype was analyzed. One pancreatic section from each mouse was selected, and three 500 µm × 500 µm regions of interest (ROI) were identified for ablation based on review of hematoxylin and eosin (H&E) stained pancreatic sections. Data collection and analysis were performed by the ImmunoMonitoring Core at the Houston Methodist Research Institute. Antibodies were conjugated to metal isotope according to the Fluidigm protocol and tissues were stained as previously described^29^. Briefly, the tissue sections were baked at 60 °C overnight, followed by deparaffinization in xylene and rehydration in alcohol gradients for 10 min each. Epitope retrieval was performed in a water bath at 95 °C in Tris-Tween 20 buffer (pH 9) for 20 min. Sections were blocked with 3% BSA in tris-buffered saline, followed by overnight staining at 4 °C with the panel of 22 markers and subsequent nuclear staining with Cell-ID Intercalator (Standard BioTools). Slides were air-dried and ablated using Hyperion Imaging System (Standard BioTools) for data acquisition.

The IMC data were preprocessed and checked for tissue integrity, staining quality, and signal range prior to analysis using customized Python scripts^30^. For every ROI, single cells were segmented based on DNA staining (Ir191) and other cell surface markers^31^. Following cell segmentation, data were processed using the Histology Topography Cytometry Analysis Toolbox (HistoCAT), and the mean intensity of each marker was extracted for each cell. Data from all samples were consolidated using R scripts for downstream analysis. The intensity values for each marker were clipped at the 99.5th percentile to eliminate outliers and subsequently normalized for unsupervised clustering in Seurat using Louvain algorithm^29,32–34^. Cell clusters were annotated based on the mean expression of markers and consolidated into 17 known cell types. Cell densities of each cell type were calculated by normalizing cell counts to the corresponding ROI area. Neighborhood analysis identified the statistically significant neighboring status for each pair of cell types^32^. The neighborhood heatmap normalized the results to -1 to 1, where 1 (or red) denoted two cell types are neighboring each other, -1 (or blue) denoted they are significantly avoiding each other, and 0 indicated not significant spatial relationship is observed between them. Customized R scripts were used for visualizing single cell clusters, neighborhood analyses, and spatially mapping the annotated cell types.

### Primary Pancreatic ADM Culture

C57BL/6 WT and CD73^-/-^ mice (10-12 weeks old) were euthanized and their pancreata were harvested. Pancreata were washed in separate tubes with cold 1X HBSS. The tissue was minced into 1 to 3 mm^3^ pieces in a petri dish, washed with HBSS and then centrifuged for 2 minutes at 450 x g at 4°C. The supernatant was discarded to remove cell fragments and blood cells. Collagenase IA solution (1X HBSS containing 10 mM HEPES, 200 U/ml of collagenase IA, and 0.25 mg/ml of trypsin inhibitor) was added and the mixture was incubated at 37 °C for 30 min. During this time, mechanical dissociation was performed by vigorously pipetting the tissue approximately 10 times using a sterile 5-mL serological pipette. Buffered washing solution (1X HBSS 1x containing 5 % FBS and 10 mM HEPES) was added to stop the digestion. Tubes were centrifuged for 3 minutes at 450 x g at 4°C and the supernatant was aspirated. Pellets were resuspended in Modified Waymouth’s medium (dexamethasone 20 µg/mL, 10% FBS, 0.1 mg/ml Soybean Trypsin Inhibitor, penicillin 100 I.U./mL, and streptomycin 100µg/ml) and passed through 100 μm filters. Filters were rinsed with Modified Waymouth’s medium. The cell suspension was mixed with growth factor-reduced Matrigel at a 1:1 ratio and seeded at a density of 800,000 cells per well in a 24-well plate. The plate was incubated at 37°C for 1 hour. Next, 500 μl Waymouth’s medium was added to each well. After 24 hours, ADM began to form, and the medium was replenished every other day. Cells were cultured for three days. Culture supernatants were collected and cells were harvested on day 3^35,36^. Two mice were used per group.

### Cytokine Array and ImageJ Analysis

Proteome Profiler Mouse Cytokine Array Kit, Panel A, was purchased from R&D Systems (ARY006). Reagents were prepared according to the manufacture’s protocol. Supernatant from CD73^-/-^ and WT primary ADM culture were applied to the membranes. Membranes were incubated in a 4-well multidish with Array Buffer 6 for one hour on a rocking platform at RT. Samples were diluted in Array Buffer 4 and 6 and then added to Mouse Cytokine Array Panel A Detection Antibody Cocktail. Membranes were incubated overnight at 4^°^C on rocking platform. Membranes were separated into individual containers and washed three times with 1X Wash Buffer for 10 min per wash at RT on a rocking platform. Streptavidin-HRP was diluted in Array Buffer 6 and added to 4-well multi-dish. Membranes were incubated in Streptavidin-HRP for 30 min on a shaker. Membranes were covered with a plastic sheet protector, and ECL reagent was added to each membrane. Membranes were imaged using ChemiDoc system.

### Statistical Analysis

Statistical analyses were performed using GraphPad Prism v11.1.0. Comparisons between two groups were performed using unpaired two-tailed t-tests with Welch’s correction. For cluster density analysis, statistical comparisons were performed using two-way ANOVA followed by Sidak’s multiple comparisons test. Data are presented as mean ± SD, and P < 0.05 was considered statistically significant.

## RESULTS

### Chronic injury drives CD73+ acinar-to-ductal metaplasia

To investigate the mechanisms by which CD73 facilitates ADM resolution or progression, we first asked whether metaplasia-specific expression of CD73 is important for chronic pancreatitis pathogenesis. To determine whether CD73 is expressed in injury-induced ADM, we analyzed our previously published scRNA-seq dataset of acinar lineage-traced ADM from Ptf1aCre^ERTM/+^;LSL-Rosa^eYFP/+^ mice. Nt5e was minimally expressed in Prss1+ acinar cells, but enriched in Hnf1β+, Krt19+ ADM (Fig. 1A-B). To validate CD73 expression in ADM at the protein level, we collected pancreata from acinar lineage trace mice and performed co-immunofluorescence. As shown in Figure 1C, CD73 expression was significantly increased in YFP+CK8/18+ ADM following caerulein-induced pancreatic injury, while little to no CD73 was detected in YFP+ acinar cells in control mice. Quantification showed a higher proportion of YFP+CD73+ cells and greater CD73 fluorescence intensity in caerulein-treated pancreas compared with PBS controls (Fig. 1D). We also analyzed organoids generated from Ptf1aCre^ERTM/+^;LSL-Rosa^eYFP/+^ mice treated with either caerulein or control. YFP expression was retained in caerulein-derived ADM organoids, confirming their acinar-lineage origin (Fig. 1E). Nt5e expression was significantly higher in caerulein-derived ADM organoids compared with ductal organoids derived from PBS-treated mice (Fig. 1F). Altogether, these findings demonstrate upregulation of CD73 in acinar-derived ADM resulting from chronic pancreatic injury. CD73 is a critical checkpoint for tissue recovery in CP IHC staining of caerulein-induced murine CP demonstrated strong CD73 expression localized to ADM structures (Fig. 2A). CD73 expression was also detected in pancreatic tissues from patients with chronic pancreatitis. In one patient, CD73 expression was more clearly associated with metaplastic and ductal structures, whereas in a second patient, CD73 staining was more broadly distributed throughout the tissue (Fig. 2A) consistent with expression in both the epithelium and stroma. To determine the role of ADM- specific CD73 during chronic pancreatitis, we crossed Hnf1bCreERT2 mice with CD73^fl/fl^ mice to generate CD73^fl/fl^;Hnf1bCreERT2 mice. Control Hnf1bCreERT2 (n=7) and CD73^fl/fl^;Hnf1bCreERT2 mice (n=5) were treated with caerulein twice daily for 4 weeks to induce chronic pancreatic injury and establish ADM, followed by tamoxifen administration to delete CD73 in metaplastic epithelial cells. After 1 week of recovery, pancreas was collected for histological analysis (Fig. 2B). CD73 staining confirmed a significant reduction in epithelial-specific CD73 expression in CD73^fl/fl^;Hnf1bCreERT2 mice, while CD73 expression in blood vessels remained unchanged (Fig. 2C, D). Histological analysis showed significantly greater inter- acinar edema in CD73^fl/fl^;Hnf1bCreERT2 mice compared with control mice (Fig. 2C, E). CD73^fl/fl^;Hnf1bCreERT2 mice also showed increased collagen deposition and αSMA+ area compared with control mice (Fig. 2C, F, G), consistent with greater fibrosis. These findings suggest that metaplastic epithelial CD73 aids in the resolution of pancreatic injury and fibrosis during recovery from chronic pancreatitis.

**Figure 1.**
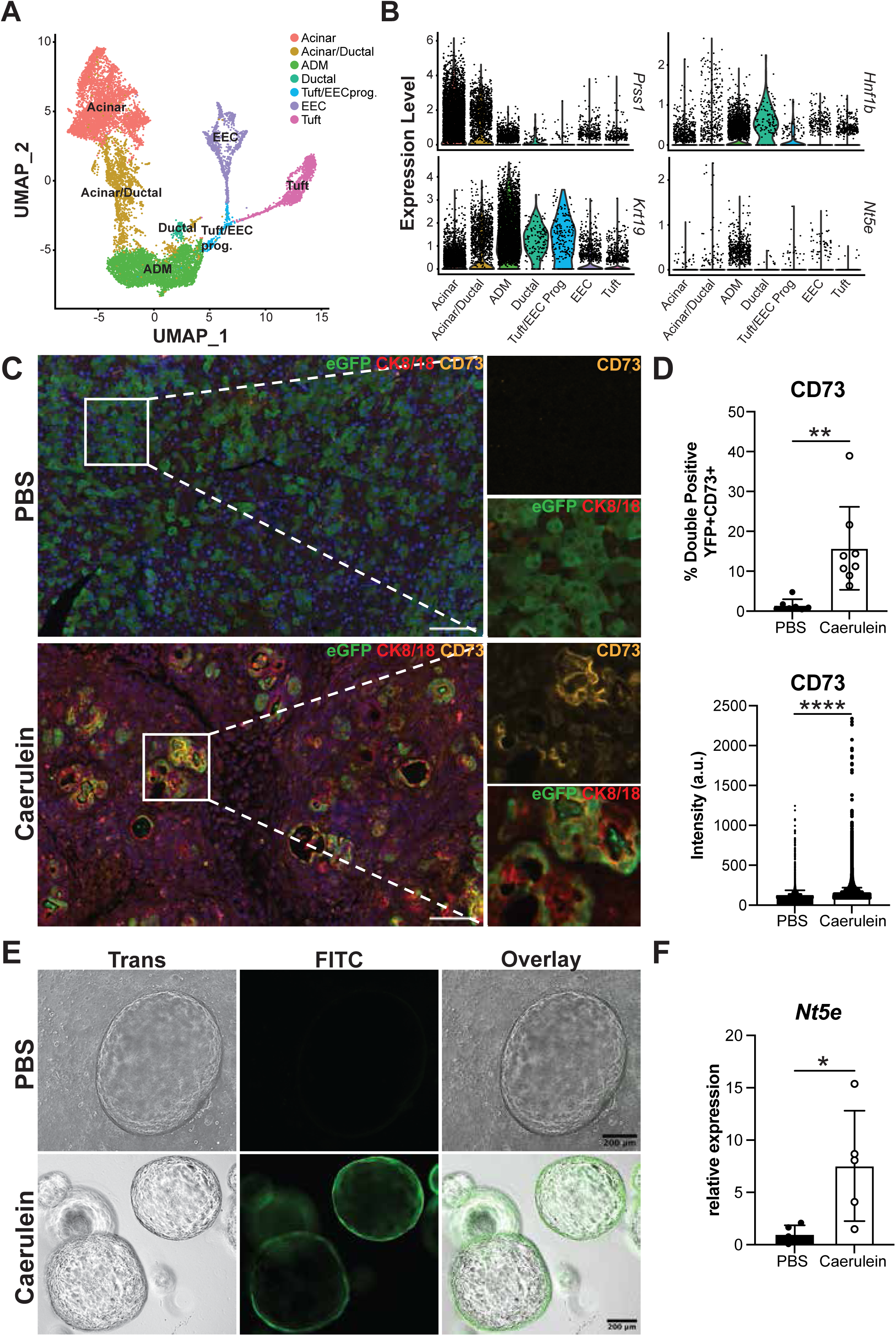
Chronic tissue injury drives CD73+ acinar to ductal metaplasia. A) Uniform manifold approximation and projection (UMAP) and B) Violin plots of lineage-traced single cell RNA-sequencing data showing expression of acinar cell marker Prss1, ductal cell markers Krt19 and Hnf1b, and Nt5e across metaplastic populations in chronic pancreatitis derived from Ptf1aCre^ERTM/+^;LSL-Rosa^eYFP/+^ mice treated with caerulein. B) Co-immunofluorescence for GFP (lineage trace; green), CD73 (yellow), and Krt8/18 (ductal/ADM marker; red) in pancreata from the lineage trace model treated with either caerulein or control PBS, quantified in C)(n = 8 caerulein; n = 6 PBS) Scale bars, 50μm. D) Brightfield or FITC imaging of organoids generated from pancreata from the lineage trace model treated with either caerulein or PBS. Scale bars, 200μm. E) qRT-PCR for Nt5e/CD73 in normal or ADM organoids (n = 5 caerulein; n = 4 PBS). Data are presented as mean ± SD. Statistical significance was determined using an unpaired t test with Welch’s correction. *, p < 0.05; **, p < 0.01; ****, p < 0.001.

**Figure 2.**
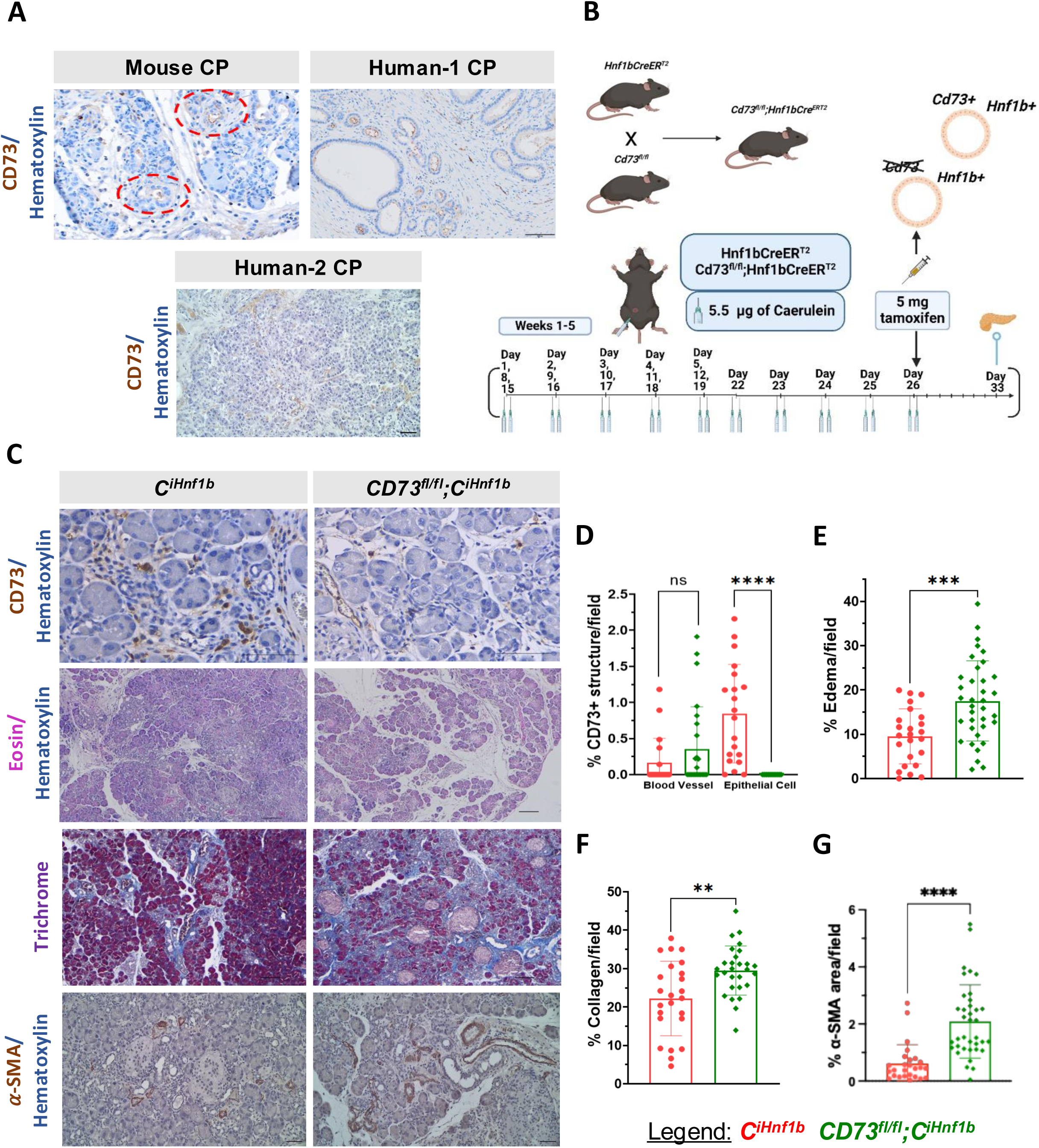
CD73 is a critical checkpoint for tissue recovery in CP. A) Representative images of pancreatic tissues from a mouse with chronic pancreatitis and two patients with chronic pancreatitis showing CD73 expression. B) Schematic illustrating the generation of CD73^fl/fl^;Hnf1bCreERT2 mice by crossing Hnf1bCreERT2 mice with CD73^fl/fl^ mice and the experimental study design. Hnf1bCreERT2 control (n=7) and CD73^fl/fl^;Hnf1bCreERT2 (n=5) mice received 5.5 μg caerulein for 4 weeks. On day 26, all mice received 5 mg tamoxifen. Mice were euthanized on day 33, and pancreas was collected for analysis. C) Representative images of pancreatic tissues from Hnf1bCreERT2 and CD73^fl/fl^;Hnf1bCreERT2 mice showing CD73 expression, H&E staining, trichrome staining and α-smooth muscle actin (α-SMA) expression. Scale bars, 50 μm. Quantification of D) % CD73+ regions within blood vessels and epithelial cells, E) % edema area, F) % collagen+ area and G) % αSMA+ area per field in Hnf1bCreERT2 and CD73^fl/fl^;Hnf1bCreERT2 mice. Quantifications were performed for n = 3–5 mice per group and 5–10 fields analyzed per mouse. Data are presented as mean ± SD. Statistical significance was determined using an unpaired t test with Welch’s correction. *p<0.05, **p<0.01, ***p<0.001, ****p<0.0001.

Loss of metaplastic Cd73 impairs epithelial proliferation and promotes persistent ADM in CP

We next examined whether loss of metaplastic CD73 affects acinar regeneration during recovery from chronic pancreatitis. Pancreata from CD73^fl/fl^;Hnf1bCreERT2 mice showed reduced proliferative activity, as indicated by fewer PCNA+ nuclei and Ki67+ cells per field compared with control Hnf1b-CreERT2 mice (Fig. 3A–C). In addition, CD73^fl/fl^;Hnf1bCreERT2 pancreata showed a significantly lower percentage of α- amylase+ area and a higher percentage of CK19+ ductal area than controls (Fig. 3A, D, E), suggesting impaired recovery of the acinar compartment and persistence of metaplastic epithelial structures. These findings suggest that metaplastic CD73 supports acinar regeneration and restoration of normal pancreatic architecture during recovery from chronic pancreatitis.

**Figure 3.**
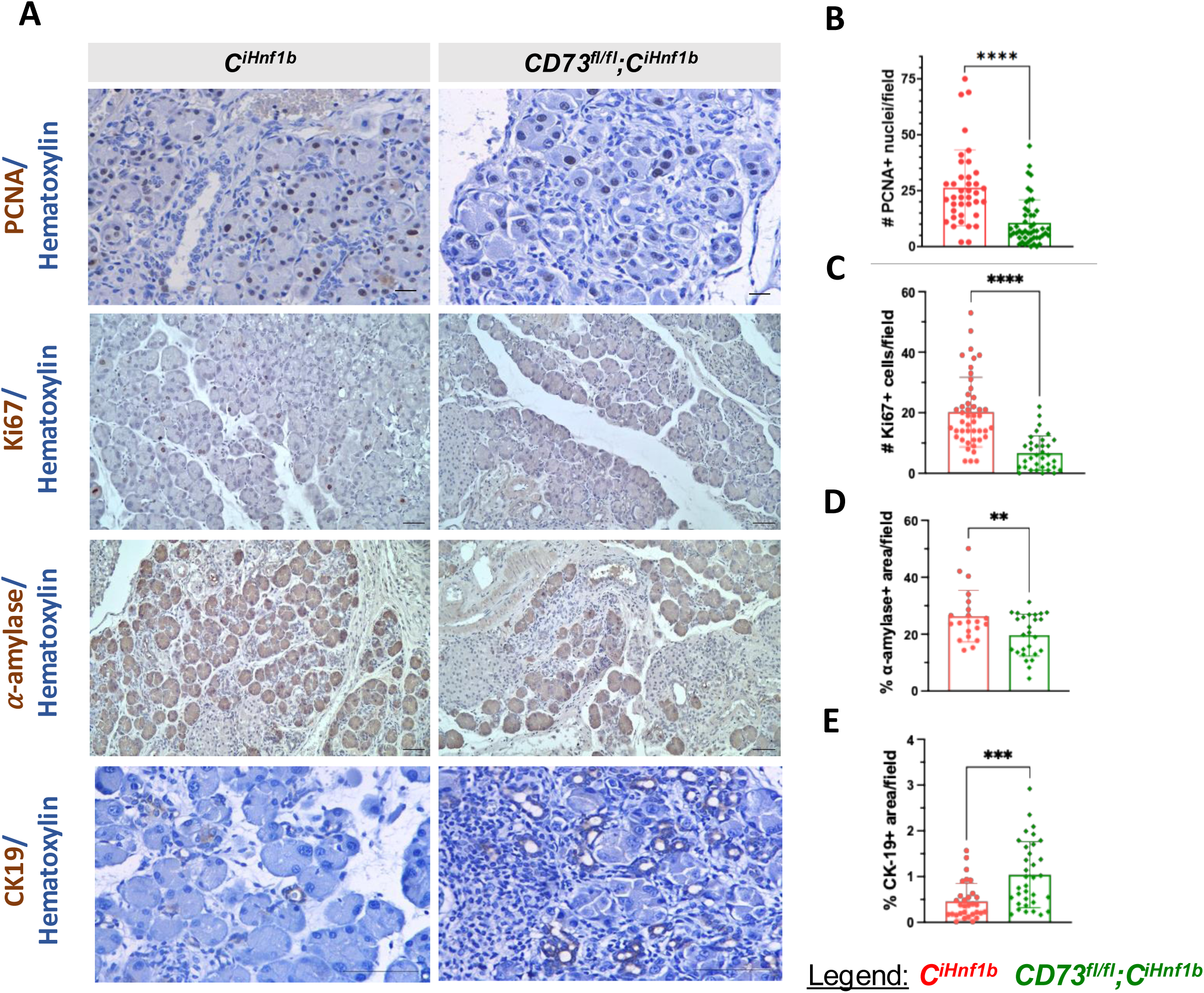
CD73 regulates epithelial plasticity during CP. A) Representative images of pancreatic tissues from Hnf1bCreERT2 and CD73^fl/fl^;Hnf1bCreERT2 mice showing PCNA, Ki67, α-amylase and CK19 expression. Scale bars, 50 μm. Quantification of B) PCNA+ nuclei, C) Ki67+ cells, D) % α-amylase+ area and E) % CK19+area per field in Hnf1bCreERT2 and CD73^fl/fl^;Hnf1bCreERT2 mice. Quantifications were performed for n = 3–5 mice per group and 5–10 fields analyzed per mouse. Data are presented as mean ± SD. Statistical significance was determined using an unpaired test with Welch’s correction. *p<0.05, **p<0.01, ***p<0.001, ****p<0.0001.

Metaplasia specific CD73 deletion alters inflammatory cell composition and spatial organization To explore how loss of metaplastic CD73 affects the inflammatory microenvironment during recovery from CP, we performed spatial immune profiling using IMC. Overall, eighteen clusters representing immune, epithelial, endothelial and stromal populations were identified (Fig. 4A-B). Loss of ADM-specific CD73 in CD73^fl/fl^;Hnf1bCreERT2 mice resulted in higher cluster densities of NK cells and Ly6G⁺CD44- tissue resident neutrophils as compared to Hnf1bCreERT2 pancreata with intact CD73. Conversely, CD4⁺ T cells were more abundant in ADM-CD73+ tissues (Fig. 4C), suggesting that CD73 loss skews immune composition toward increased or persistent innate inflammatory cell infiltration. Further, CD73-floxed tissue showed increased MHC II+ cluster density, whereas CD8α+, Granzyme B (Gzmb)+, and Foxp3+ cluster densities were not significantly different between groups (Supplementary Fig. 1A-B). To characterize immune infiltration in a spatial manner, we performed neighborhood analysis of the IMC data. In Hnf1bCreERT2 pancreata, Ly6G⁺CD44- neutrophils localized adjacent to endomucin+ endothelial cells, whereas in CD73fl/fl;Hnf1bCreERT2 mice, neutrophils instead shifted toward PanCK⁺αSMA⁺ ADM, indicating loss of neutrophil association with vasculature and increased association with metaplastic epithelial structures following CD73 deletion. Additionally, in CD73^fl/fl^;Hnf1bCreERT2 mice, neutrophils were localized near B cells, CD11c+ dendritic cells and Ly6C+ monocytes. CD11c+ dendritic cells likewise shifted tissue compartments. In Hnf1bCreERT2 pancreata, dendritic cells localized near ADM; in metaplasia CD73-deficient pancreata, they preferentially associated with endothelial structures, suggesting impaired infiltration. F4/80+ macrophages remained distributed near both ADM and the vasculature regardless of genotype (Fig. 4D-E).

**Figure 4.**
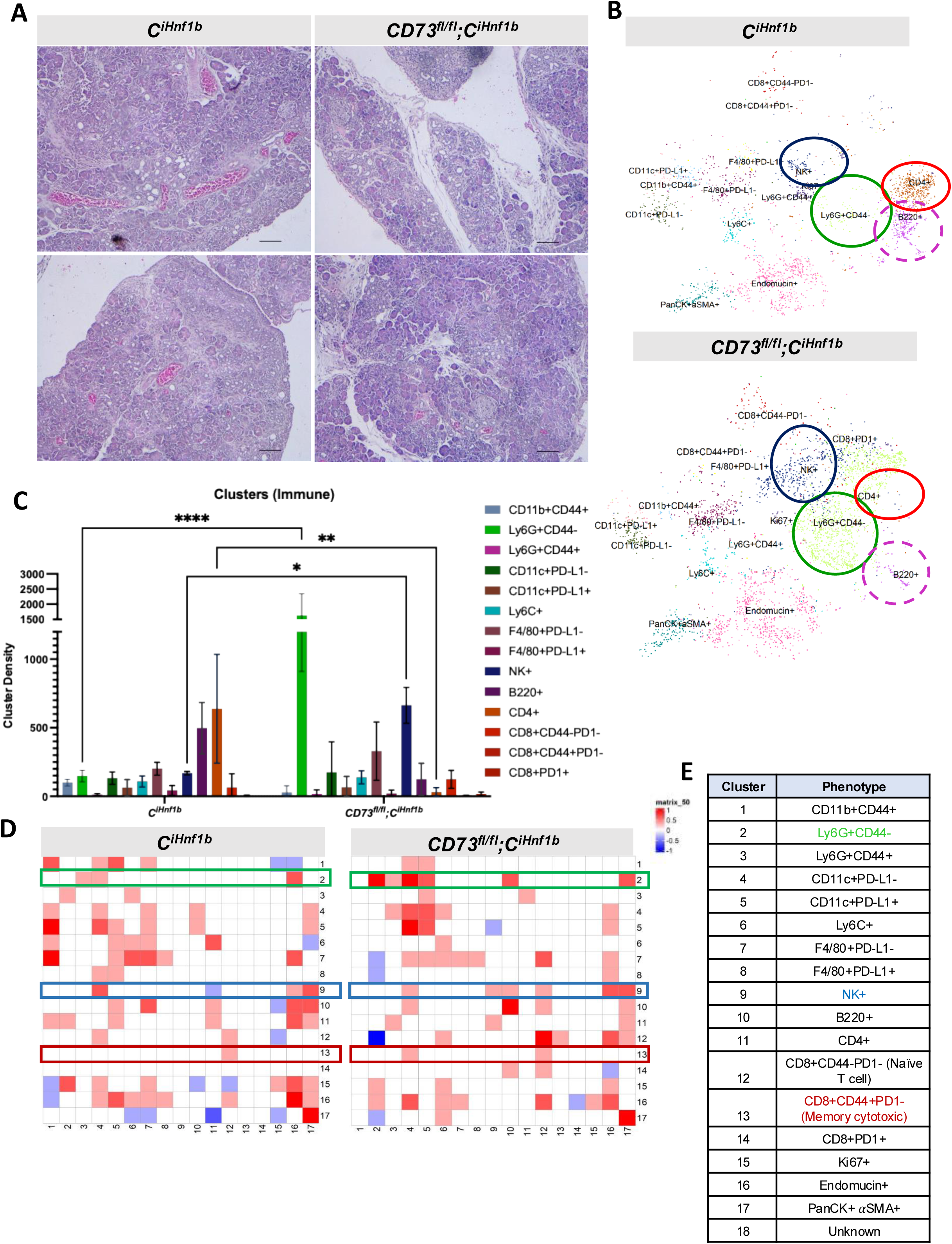
Loss of ADM CD73 alters the spatial organization of immune cells in the CP microenvironment. A) Representative images of pancreatic tissues showing the regions selected for imaging mass cytometry analysis. B) Cluster-density analysis showing the immune-cell profiles of pancreatic tissues from Hnf1bCreERT2 (n=1) and CD73^fl/fl^;Hnf1bCreERT2 (n=1) mice. C) Quantification of the densities of NK+, Ly6G+CD44−, B220+ and CD4+ cell phenotypes in pancreatic tissues from Hnf1bCreERT2 and CD73^fl/fl^;Hnf1bCreERT2 mice. D) Neighborhood analysis of IMC data presented as a heatmap. Red indicates cell populations that are spatially associated linked or located in proximity, whereas blue indicates cell populations that are spatially separated or less likely to be located near one another. E) Table showing the cell cluster numbers and their corresponding cellular phenotypes used in the neighborhood analysis heatmaps in panel D.

Although NK cell abundance increased overall in ADM-CD73 deficient tissues, NK+ clusters localized near both ADM and endothelial regions in both genotypes. Activated memory CD8+CD44+ T cells were observed adjacent to ADM and endothelium only in CD73-deficient tissue, suggesting enhanced T cell activation in the absence of CD73 (Fig. 4D-E). Collectively, these exploratory data demonstrate that epithelial CD73 may influence both immune-cell composition and spatial organization during CP. Metaplasia deficient in CD73 promote pro-inflammatory immune cell abundance in CP Consistent with the IMC findings, CD73^fl/fl^;Hnf1bCreERT2 mice showed a significantly higher percentage of NIMPR14+ neutrophils compared with control mice, whereas the percentage of MPO+ myeloid cell area was unchanged between the two groups (Fig. 5A–C). Additionally, the percentage of CD68+ monocytes/macrophages, the number of CD163+ cells, and the percentage of CCR2+ inflammatory monocytes per field were significantly reduced in CD73^fl/fl^;Hnf1bCreERT2 mice (Fig. 5A, D–F). The number of NK1.1+ natural killer cells was significantly increased in CD73^fl/fl^;Hnf1bCreERT2 mice compared with controls (Fig. 5A, G). Furthermore, TIMP-1, a regulator of extracellular matrix remodeling, has been reported to be elevated in patients with chronic pancreatitis^37^. CD73^fl/fl^;Hnf1bCreERT2 mice also showed a higher percentage of TIMP-1+ area than controls (Fig. 5A, H). Together, these findings indicate that loss of metaplastic CD73 alters the innate immune response during recovery from chronic pancreatitis, with reduced macrophage and CCR2+ cell populations, increased NK1.1+ cells, and enhanced TIMP-1 expression.

**Figure 5.**
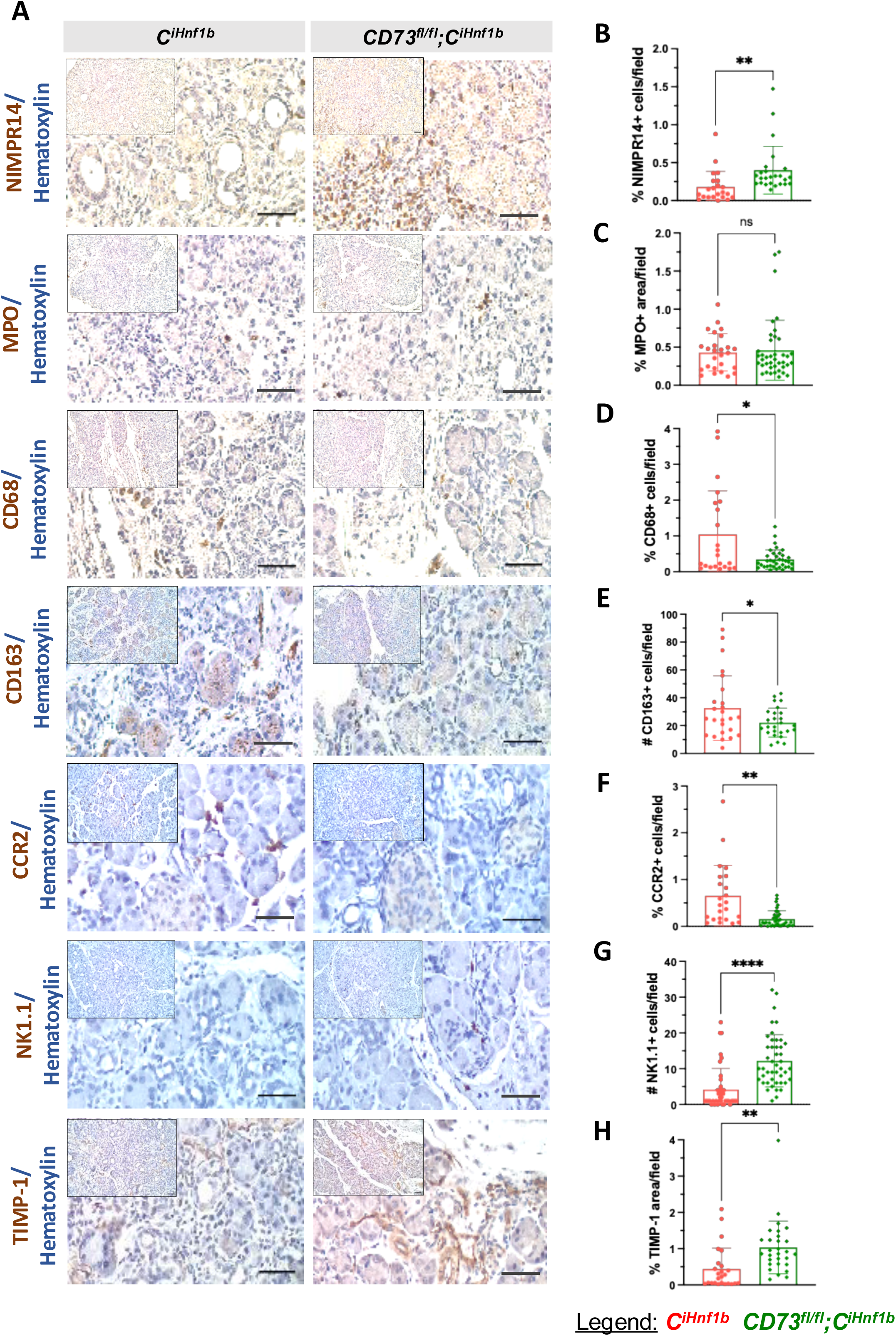
Loss of ADM CD73 results in significant NK cell and neutrophil infiltration. A) Representative images of pancreatic tissues from Hnf1bCreERT2 and CD73^fl/fl^;Hnf1bCreERT2 mice showing NIMPR14, MPO, CD68, CD163, CCR2, NK1.1 and TIMP-1 expression. Scale bars, 50 μm. Quantification of B) %NIMPR14+ cells, C) % MPO+ area, D) % CD68+ cells, E) CD163+ cells, F) % CCR2+ cells, G) NK1.1+ cells and H) % TIMP-1+ area per field in Hnf1b-CreERT2 and CD73^fl/fl^;Hnf1bCreERT2 mice. Quantifications were performed for n = 3–5 mice per group and 5–10 fields analyzed per mouse. Data are presented as mean ± SD. Statistical significance was determined using an unpaired t test with Welch’s correction. *p<0.05, **p<0.01, ***p<0.001, ****p<0.0001.

CD73 deficiency promotes ex vivo ADM formation and inflammatory mediator production To determine whether the loss of epithelial CD73 impacts ADM and inflammation in a cell autonomous manner, we employed an ex vivo model in which primary acinar cells are isolated from mice, then embedded and cultured in Matrigel. We isolated normal acini from either wild type or full-body CD73^⁻/⁻^ mice and allowed them to undergo ADM (Fig. 6A). Morphologic assessment revealed that, consistent with in vivo data in Fig. 3, CD73^-/-^ cultures generated significantly more ADM structures at both Days 1 and 3 as compared to wild type cultures (Fig. 6B-C). Supernatants collected from Day 3 ADM cultures showed significantly elevated MCP-1 (CCL2) and TIMP-1 in CD73⁻/⁻ cultures by proteome profiler array analysis (Fig. 6D), consistent with the increased TIMP-1 observed in pancreatic tissue (Fig. 3) and indicating increased inflammatory mediator production. Altogether, these data show that loss of CD73 enhances epithelial plasticity and promotes ADM formation in the absence of direct stromal or immune-cell interactions.

**Figure 6.**
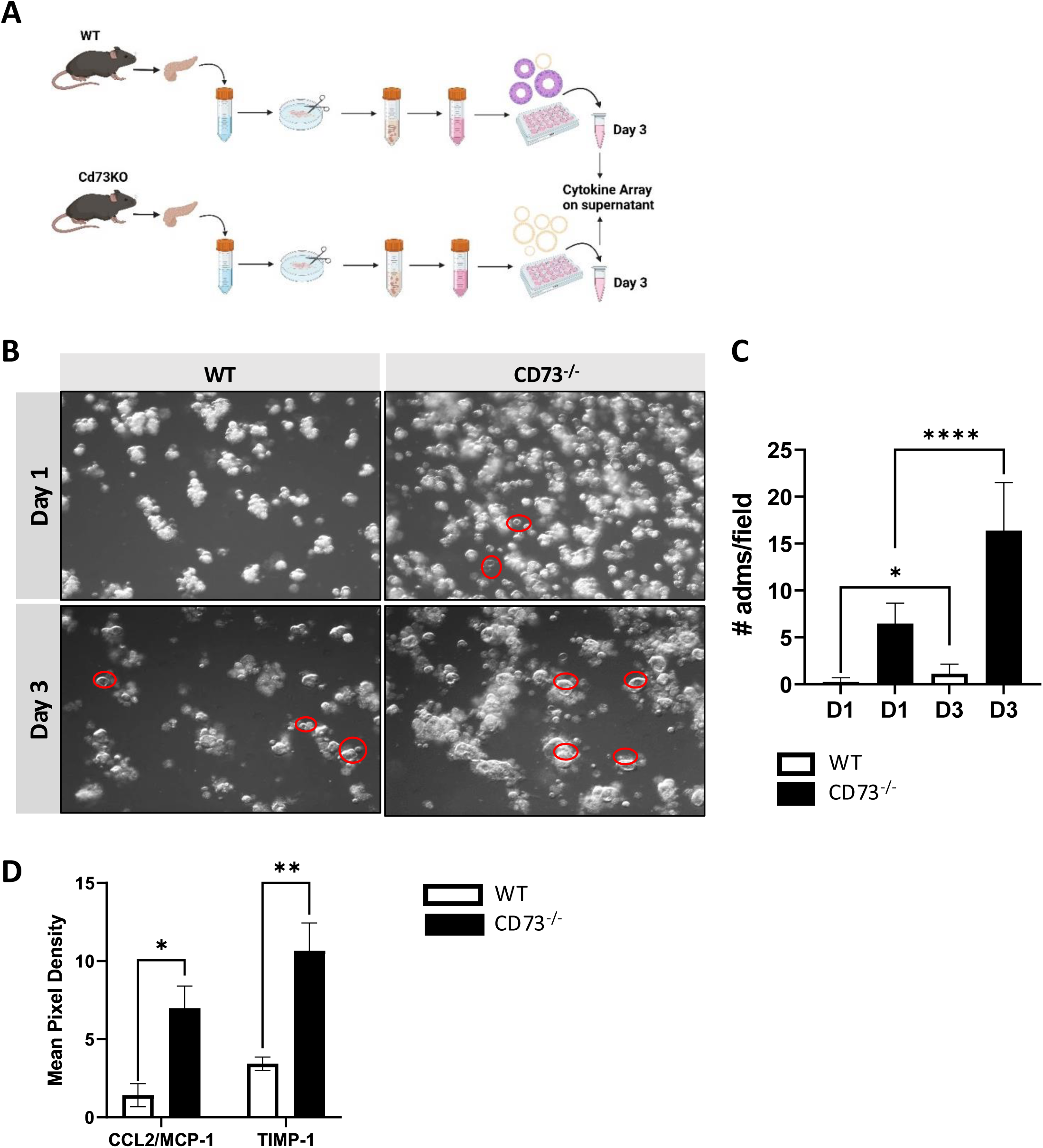
CD73-deficient acinar cells generate significantly more ADM and pro-inflammatory factor secretion in an ex vivo culture model. A) Schematic illustrating the experimental design for the ex vivo ADM assay. B) Representative 3D images of WT vs CD73-/- ADM ex vivo culture. Acinar cells were isolated and grown in Matrigel supplemented with media for three days. Red circles indicate ADM structures. C) Quantification of ADM structures formed on days 1 and 3 in cultures derived from WT and CD73-/- mice. D) Quantification of the mean pixel intensity of CCL2/MCP-1 and TIMP-1 on day 3 in cultures derived from WT and V mice. Data are presented as mean ± SD. Statistical significance was determined using two-way ANOVA. *p<0.05, **p<0.01, ***p<0.001, ****p<0.0001

## DISCUSSION

ADM represents a key epithelial plasticity program activated during pancreatic injury and is prominent in human chronic pancreatitis. Although ADM is initially reversible and contributes to pancreatic tissue recovery, persistent inflammation and oncogenic KRAS mutations can promote sustained ADM and progression toward neoplasia. Despite the recognized importance of ADM in CP and early pancreatic tumorigenesis, the epithelial factors that regulate ADM resolution and shape the local immune microenvironment remain poorly defined. Here, we identify metaplastic epithelial CD73 as an important regulator of pancreatic recovery following chronic injury. Lineage tracing in Ptf1aCre^ERTM/+^;LSL-Rosa^eYFP/+^ mice demonstrated increased CD73 expression in ADM following pancreatic injury. Further, an inducible Hnf1bCreERT2 model was used in which tamoxifen was administered after 4 weeks of caerulein treatment, when ADM was already established, allowing CD73 to be deleted specifically within HNF1β-expressing metaplastic epithelial cells. Loss of epithelial CD73 impaired epithelial proliferation and recovery of the acinar cells, increased persistent ADM and fibrosis, and altered the abundance and spatial organization of inflammatory cells. Together, these findings suggest that metaplastic epithelium orchestrates tissue recovery and immune remodeling through CD73 during chronic pancreatitis.

Our previous work demonstrated that CD73 regulates tissue recovery and inflammatory resolution during pancreatic injury. Using global CD73-deficient mice in caerulein-induced acute and chronic pancreatitis, we found that loss of CD73 delayed tissue recovery and resulted in sustained metaplasia and inflammation compared with wild type mice. CD73-deficient pancreata showed increased neutrophil accumulation, and depletion of neutrophils reduced both metaplasia and pancreatic injury, demonstrating that persistent neutrophilic inflammation contributed to impaired resolution^15^. However, because CD73 was globally deleted in that study, the contributions of individual epithelial, immune, stromal and endothelial compartments could not be distinguished.

The present study shows that loss of epithelial CD73 reduced PCNA and Ki67+ cells, decreased α- amylase+ area, and increased CK19+ area, indicating impaired epithelial proliferation, incomplete recovery of the acinar compartment, and persistent ADM. Increased edema, collagen deposition, and α- SMA staining in CD73 deficient pancreata further suggest that impaired epithelial recovery occurs together with persistent stromal activation and fibrotic remodeling.

Loss of metaplastic epithelial CD73 also altered the innate immune landscape of the chronically injured pancreas. CD73-deficient pancreata showed increased neutrophils and NK cells. The increase in neutrophils is consistent with our previous global CD73-deficient model, in which persistent neutrophilic inflammation contributed to sustained pancreatic injury and metaplasia^15^. Neutrophils have also been implicated in the progression of chronic pancreatitis, as STAT5-dependent GM-CSF production was shown to increase neutrophil infiltration and promote pancreatic inflammation and fibrosis^38^. Although NK cells can infiltrate the injured pancreas and regulate inflammatory responses, their function during chronic pancreatitis and ADM remains poorly understood. Whether the sustained metaplasia observed after CD73 loss is also mediated by NK cells requires further investigation.

In contrast, CD73-deficient pancreata showed reduced macrophage abundance. Previous studies have shown that inflammatory macrophages are more prominent during the early phase of pancreatitis, whereas macrophages involved in tissue repair become more prominent during recovery. Depletion of macrophages during the recovery phase delayed inflammatory resolution, highlighting their importance in pancreatic repair^39^. Macrophages can therefore have context-dependent functions. Inflammatory macrophage-derived signals may promote ADM, whereas other macrophage populations support the clearance of damaged tissue, acinar redifferentiation, and pancreatic recovery^6,40^. Accordingly, the reduction in CD68⁺ and CD163⁺ macrophages in our study may indicate disruption of macrophage responses required for inflammatory resolution and tissue recovery. The accompanying reduction in CCR2+ area further suggests altered monocyte–macrophage responses, although the mechanisms underlying their reduced abundance remain to be determined.

Beyond changes in immune-cell abundance, exploratory IMC and neighborhood analyses suggested that loss of metaplastic epithelial CD73 altered the spatial organization of inflammatory cells within the chronically injured pancreas. In CD73-deficient tissue, NK cells and Ly6G⁺ neutrophils were located near metaplastic epithelial structures, while the association of neutrophils with endothelial regions was reduced. These findings suggest that epithelial CD73 regulates not only the composition but also the spatial arrangement of the inflammatory microenvironment. However, because the IMC analysis was exploratory, these spatial relationships will need to be confirmed in additional biological samples.

The ex vivo findings suggest that CD73-deficient epithelial cells may influence the surrounding inflammatory and stromal environment through secreted mediators. CD73-deficient acinar cultures formed more ADM structures and released higher levels of MCP-1 and TIMP-1. Although this model does not reproduce chronic pancreatitis, the increase in TIMP-1 was consistent with its elevated expression in CD73-deficient pancreatic tissue, supporting metaplastic epithelial cells as a potential source of TIMP-1 during pancreatic injury. Recent work in pancreatic cancer identified epithelial-derived TIMP-1 as a major component of an immunoregulatory secretory program that directly suppressed NK-cell cytotoxicity and IL-2–mTOR signaling^41^. Although NK-cell function was not examined in our study, the concurrent increase in TIMP-1 and NK-cell abundance raises the possibility that the CD73-deficient metaplastic epithelium may influence not only NK-cell localization but also their functional state. MCP-1 regulates monocyte– macrophage migration through CCR2; however, increased MCP-1 production in the ex vivo ADM cultures did not correspond with increased CCR2+ area in CD73-deficient pancreatic tissue. This difference may reflect altered CCR2 expression or changes in monocyte–macrophage responses following epithelial CD73 deletion.

Our findings also raise important questions for future investigation. Defining how epithelial CD73- generated adenosine regulates immune-cell abundance and localization may clarify the mechanisms linking metaplastic epithelium with inflammatory resolution during chronic pancreatic injury. Further studies are also needed to determine how epithelial-derived MCP-1 and TIMP-1 influence monocyte– macrophage responses, NK-cell function, ADM persistence, and fibrotic remodeling. Because CD73 appears to support tissue recovery during pancreatitis but promotes immune suppression in established PDAC, it will be important to identify when its function shifts from tissue protective to tumor promoting. Preclinical models that follow chronic pancreatitis through early neoplastic progression may help define the disease stages at which CD73 or adenosine receptor inhibition can limit tumor-promoting immune suppression without exacerbating tissue injury in high-risk patients.

In summary, our findings identify CD73 as an important regulator linking epithelial plasticity, tissue recovery, and immune remodeling during chronic pancreatic injury. Loss of CD73 within established metaplasia impaired recovery of the acinar compartment, promoted persistent ADM and fibrosis, and altered the abundance and spatial organization of inflammatory cells. Together, these findings provide a framework for understanding how metaplastic epithelium shapes pancreatic recovery through CD73 and highlight the context-dependent role of CD73 across pancreatic injury and neoplastic disease.

---

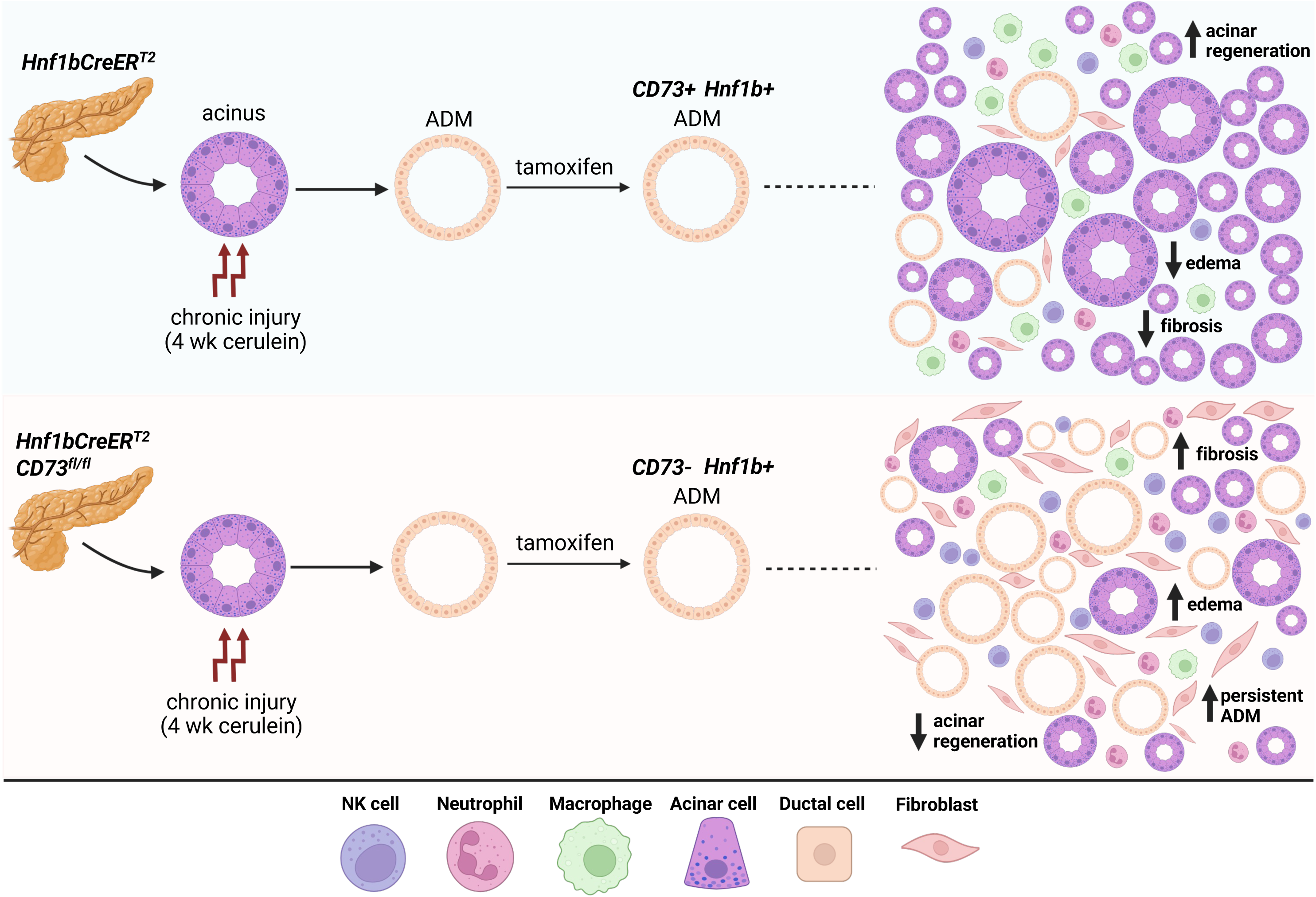

## REFERENCES

1. Beyer G, Habtezion A, Werner J, Lerch MM, Mayerle J. Chronic pancreatitis. Lancet. Aug 15 2020;396(10249):499–512. doi:10.1016/S0140-6736(20)31318-0

2. Strobel O, Dor Y, Alsina J, et al. In vivo lineage tracing defines the role of acinar-to-ductal transdifferentiation in inflammatory ductal metaplasia. Gastroenterology. Dec 2007;133(6):1999–2009. doi:10.1053/j.gastro.2007.09.009

3. Kirkegard J, Cronin-Fenton D, Heide-Jorgensen U, Mortensen FV. Acute Pancreatitis and Pancreatic Cancer Risk: A Nationwide Matched-Cohort Study in Denmark. Gastroenterology. May 2018;154(6):1729–1736. doi:10.1053/j.gastro.2018.02.011

4. Habtezion A. Inflammation in acute and chronic pancreatitis. Curr Opin Gastroenterol. Sep 2015;31(5):395–9. doi:10.1097/MOG.0000000000000195

5. Zhang X, Zhao Y, Qin C, et al. Acinar-ductal metaplasia in pancreatitis and pancreatic ductal adenocarcinoma. Cell Oncol (Dordr). Jun 29 2026;49(5)doi:10.1007/s13402-026-01251-0

6. Liou GY, Doppler H, Necela B, et al. Macrophage-secreted cytokines drive pancreatic acinar-to-ductal metaplasia through NF-kappaB and MMPs. J Cell Biol. Aug 5 2013;202(3):563–77. doi:10.1083/jcb.201301001

7. Liou GY, Fleming Martinez AK, Doppler HR, Bastea LI, Storz P. Inflammatory and alternatively activated macrophages independently induce metaplasia but cooperatively drive pancreatic precancerous lesion growth. iScience. Jun 16 2023;26(6):106820. doi:10.1016/j.isci.2023.106820

8. Zhang Y, Yan W, Mathew E, et al. Epithelial-Myeloid cell crosstalk regulates acinar cell plasticity and pancreatic remodeling in mice. Elife. Oct 5 2017;6doi:10.7554/eLife.27388

9. Murphy PS, Wang J, Bhagwat SP, et al. CD73 regulates anti-inflammatory signaling between apoptotic cells and endotoxin-conditioned tissue macrophages. Cell Death Differ. Mar 2017;24(3):559–570. doi:10.1038/cdd.2016.159

10. Synnestvedt K, Furuta GT, Comerford KM, et al. Ecto-5’-nucleotidase (CD73) regulation by hypoxia- inducible factor-1 mediates permeability changes in intestinal epithelia. J Clin Invest. Oct 2002;110(7):993–1002. doi:10.1172/JCI15337

11. Vieira C, Magalhaes-Cardoso MT, Ferreirinha F, et al. Feed-forward inhibition of CD73 and upregulation of adenosine deaminase contribute to the loss of adenosine neuromodulation in postinflammatory ileitis. Mediators Inflamm. 2014;2014:254640. doi:10.1155/2014/254640

12. Yegutkin GG, Samburski SS, Jalkanen S, Novak I. ATP-consuming and ATP-generating enzymes secreted by pancreas. J Biol Chem. Oct 6 2006;281(40):29441–7. doi:10.1074/jbc.M602480200

13. Leone RD, Emens LA. Targeting adenosine for cancer immunotherapy. J Immunother Cancer. Jun 18 2018;6(1):57. doi:10.1186/s40425-018-0360-8

14. Roh M, Wainwright DA, Wu JD, Wan Y, Zhang B. Targeting CD73 to augment cancer immunotherapy. Curr Opin Pharmacol. Aug 2020;53:66–76. doi:10.1016/j.coph.2020.07.001

15. O’Brien BJ, Faraoni EY, Strickland LN, et al. CD73-generated extracellular adenosine promotes resolution of neutrophil-mediated tissue injury and restrains metaplasia in pancreatitis. FASEB J. Jan 2023;37(1):e22684. doi:10.1096/fj.202201537R

16. Chen Q, Pu N, Yin H, et al. CD73 acts as a prognostic biomarker and promotes progression and immune escape in pancreatic cancer. J Cell Mol Med. Aug 2020;24(15):8674–8686. doi:10.1111/jcmm.15500

17. Faraoni EY, Singh K, Chandra V, et al. CD73-Dependent Adenosine Signaling through Adora2b Drives Immunosuppression in Ductal Pancreatic Cancer. Cancer Res. Apr 4 2023;83(7):1111–1127. doi:10.1158/0008-5472.CAN-22-2553

18. Ma Z, Dong M, Pan H, et al. ENPP1-Regulated Extracellular Purine Metabolism Drives Pancreatitis- Mediated Pancreatic Cancer. Gastroenterology. Apr 2026;170(4):735–752. doi:10.1053/j.gastro.2025.09.032

19. Faraoni EY, Ju C, Robson SC, Eltzschig HK, Bailey-Lundberg JM. Purinergic and Adenosinergic Signaling in Pancreatobiliary Diseases. Front Physiol. 2022;13:849258. doi:10.3389/fphys.2022.849258

20. Ma Z, Lytle NK, Chen B, et al. Single-Cell Transcriptomics Reveals a Conserved Metaplasia Program in Pancreatic Injury. Gastroenterology. Feb 2022;162(2):604–620 e20. doi:10.1053/j.gastro.2021.10.027

21. Pan FC, Bankaitis ED, Boyer D, et al. Spatiotemporal patterns of multipotentiality in Ptf1a-expressing cells during pancreas organogenesis and injury-induced facultative restoration. Development. Feb 2013;140(4):751–64. doi:10.1242/dev.090159

22. Strickland LN, Liu W, Hussein U, et al. Preventive Treatment with a CD73 Small Molecule Inhibitor Enhances Immune Surveillance in K-Ras Mutant Pancreatic Intraepithelial Neoplasia. Cancer Prev Res (Phila). Oct 1 2024;17(10):457–470. doi:10.1158/1940-6207.CAPR-24-0200

23. Sardarni UK, Faraoni EY, Waller AM, et al. Dietary Fat Content Influences PanIN Progression and Pancreatic Cancer Development in Mice. Cancer Res Commun. May 1 2026;6(5):1036–1047. doi:10.1158/2767-9764.CRC-25-0777

24. Bankhead P, Loughrey MB, Fernandez JA, et al. QuPath: Open source software for digital pathology image analysis. Sci Rep. Dec 4 2017;7(1):16878. doi:10.1038/s41598-017-17204-5

25. Goldsborough T, O’Callaghan A, Inglis F, et al. A novel channel invariant architecture for the segmentation of cells and nuclei in multiplexed images using InstanSeg. bioRxiv. 2024:2024.09.04.611150. doi:10.1101/2024.09.04.611150

26. Boj SF, Hwang CI, Baker LA, et al. Organoid models of human and mouse ductal pancreatic cancer. Cell. Jan 15 2015;160(1-2):324–38. doi:10.1016/j.cell.2014.12.021

27. Huch M, Bonfanti P, Boj SF, et al. Unlimited in vitro expansion of adult bi-potent pancreas progenitors through the Lgr5/R-spondin axis. EMBO J. Oct 16 2013;32(20):2708–21. doi:10.1038/emboj.2013.204

28. Stuart T, Butler A, Hoffman P, et al. Comprehensive Integration of Single-Cell Data. Cell. Jun 13 2019;177(7):1888–1902 e21. doi:10.1016/j.cell.2019.05.031

29. Xu Y, Zhang L, Thaiparambil J, et al. Patients with Lung Cancer of Different Racial Backgrounds Harbor Distinct Immune Cell Profiles. Cancer Res Commun. Aug 2022;2(8):884–893. doi:10.1158/2767-9764.CRC-22-0057

30. Windhager J, Zanotelli VRT, Schulz D, et al. An end-to-end workflow for multiplexed image processing and analysis. Nat Protoc. Nov 2023;18(11):3565–3613. doi:10.1038/s41596-023-00881-0

31. Greenwald NF, Miller G, Moen E, et al. Whole-cell segmentation of tissue images with human-level performance using large-scale data annotation and deep learning. Nat Biotechnol. Apr 2022;40(4):555–565. doi:10.1038/s41587-021-01094-0

32. Chung AW, Anand K, Anselme AC, et al. A phase 1/2 clinical trial of the nitric oxide synthase inhibitor L-NMMA and taxane for treating chemoresistant triple-negative breast cancer. Sci Transl Med. Dec 15 2021;13(624):eabj5070. doi:10.1126/scitranslmed.abj5070

33. Hao Y, Hao S, Andersen-Nissen E, et al. Integrated analysis of multimodal single-cell data. Cell. Jun 24 2021;184(13):3573–3587 e29. doi:10.1016/j.cell.2021.04.048

34. Schapiro D, Jackson HW, Raghuraman S, et al. histoCAT: analysis of cell phenotypes and interactions in multiplex image cytometry data. Nat Methods. Sep 2017;14(9):873–876. doi:10.1038/nmeth.4391

35. Da Silva L, Bray JK, Bulut G, Jiang J, Schmittgen TD. Method for improved integrity of RNA isolated from Matrigel cultures. MethodsX. 2020;7:100966. doi:10.1016/j.mex.2020.100966

36. da Silva L, Jiang J, Perkins C, et al. Pharmacological inhibition and reversal of pancreatic acinar ductal metaplasia. Cell Death Discov. Sep 2 2022;8(1):378. doi:10.1038/s41420-022-01165-4

37. Prokopchuk O, Grunwald B, Nitsche U, et al. Elevated systemic levels of the matrix metalloproteinase inhibitor TIMP-1 correlate with clinical markers of cachexia in patients with chronic pancreatitis and pancreatic cancer. BMC Cancer. Feb 2 2018;18(1):128. doi:10.1186/s12885-018-4055-9

38. Lin Y, Chen Y, Feng W, et al. STAT5 promotes chronic pancreatitis by enhancing GM-CSF-dependent neutrophil augmentation. J Leukoc Biol. Aug 2021;110(2):293–300. doi:10.1002/JLB.3MA1020-647R

39. Wu J, Zhang L, Shi J, et al. Macrophage phenotypic switch orchestrates the inflammation and repair/regeneration following acute pancreatitis injury. EBioMedicine. Aug 2020;58:102920. doi:10.1016/j.ebiom.2020.102920

40. Xiang H, Yu H, Zhou Q, et al. Macrophages: A rising star in immunotherapy for chronic pancreatitis. Pharmacol Res. Nov 2022;185:106508. doi:10.1016/j.phrs.2022.106508

41. Fradrich J, Reyes CM, Hendel M, et al. Multimodal profiling of pancreatic cancer reveals a TIMP-1- dominated secretory profile determining pro-tumor immunoinstruction in human cancers. Cell Rep Med. Jan 20 2026;7(1):102546. doi:10.1016/j.xcrm.2025.102546

