## Supplemental Figure 1 for "Metaplasia Orchestrates Tissue Recovery and Immune Remodeling Through CD73 in Chronic Pancreatitis"

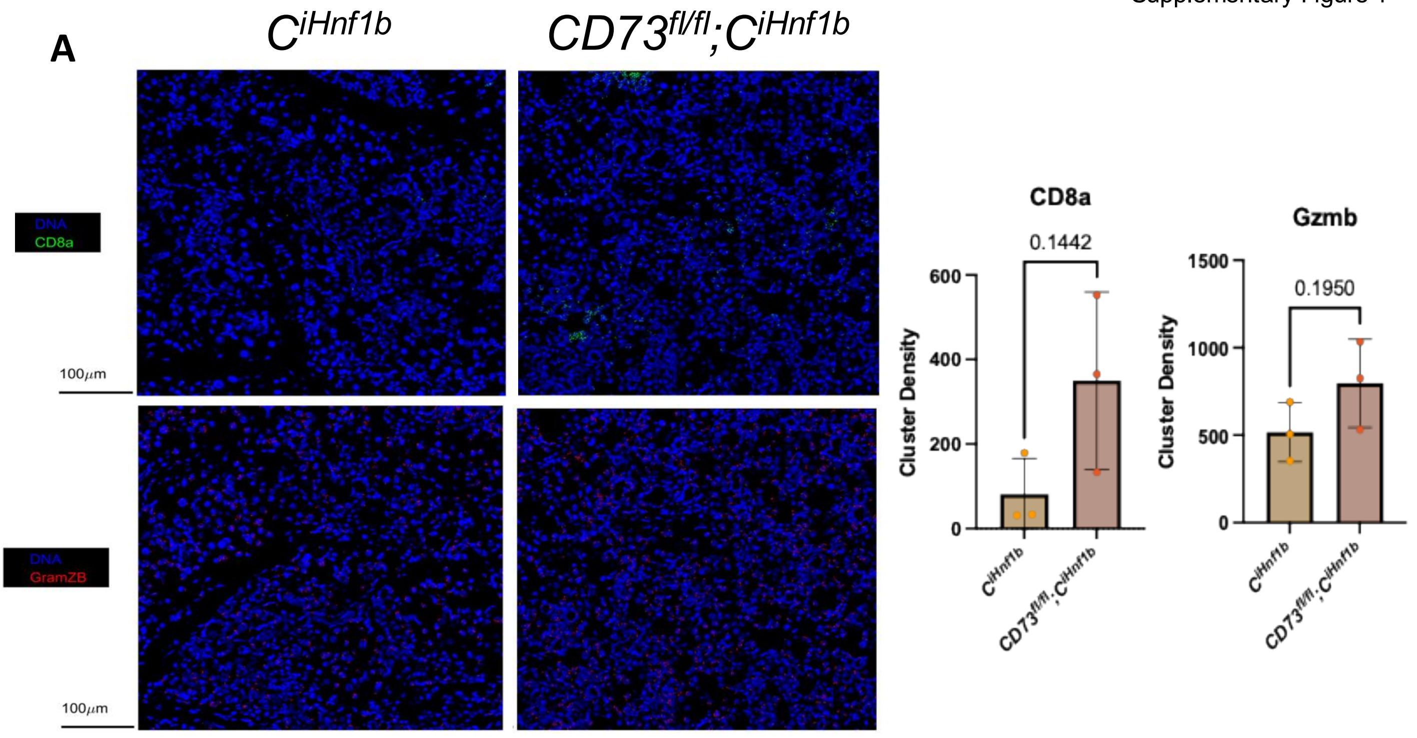


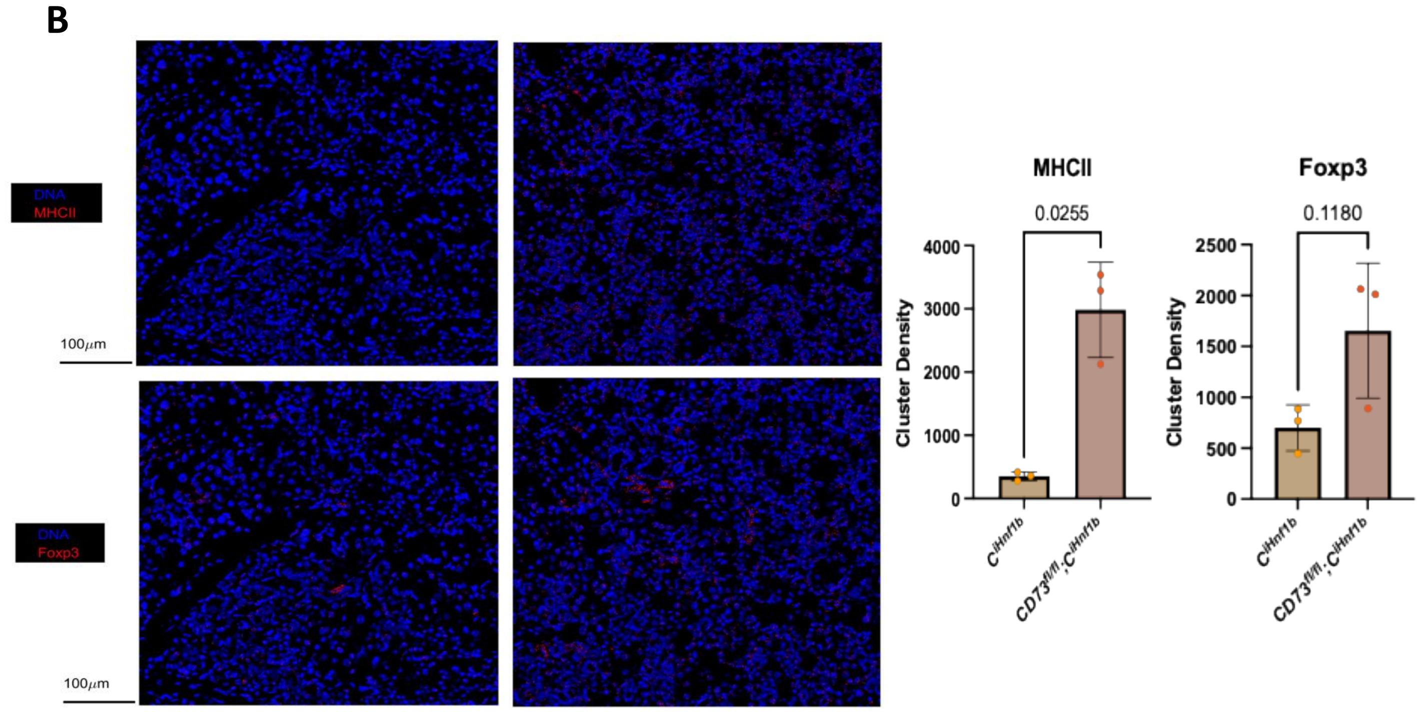


**Supplementary Figure 1. Loss of ADM CD73 is associated with increased MHC II+ cluster density. A)** Representative IMC images and quantification of the cluster densities of CD8α+ and Granzyme B (Gzmb)+ cell phenotypes in pancreatic tissues from *Hnf1bCreERT2* and *CD73^fl/fl^;Hnf1bCreERT2* mice. **B)** Representative IMC images and quantification of the cluster densities of MHC II+ and Foxp3+ cell phenotypes in pancreatic tissues from *Hnf1bCreERT2* and *CD73^fl/fl^;Hnf1bCreERT2* mice.
